# Precise annotation of *Drosophila* mitochondrial genomes leads to insights into AT-rich regions

**DOI:** 10.1101/2021.06.05.447193

**Authors:** Guangcai Liang, Jia Chang, Tung On Yau, Xin Li, Bingjun He, Jishou Ruan, Wenjun Bu, Shan Gao

**Author notes:** Corresponding authors. SG, WB. These authors contributed equally to this paper.

## Abstract

In the present study, we performed precise annotation of *Drosophila melanogaster*, *D. simulans*, *D. grimshawi*, *Bactrocera oleae* mitochondrial (mt) genomes by pan RNA-seq analysis. Our new annotations corrected or modified some of the previous annotations and two important findings were reported for the first time, including the discovery of the conserved polyA(+) and polyA(−) motifs in the control regions (CRs) of insect mt genomes and the adding of CCAs to the 3’ ends of two antisense tRNAs in *D. melanogaster* mt genome. Using PacBio cDNA-seq data from *D. simulans*, we precisely annotated the Transcription Initiation Sites (TISs) of the mt Heavy and Light strands in *Drosophila* mt genomes and reported that the polyA(+) and polyA(−) motifs in the CRs are associated with TISs. The discovery of the conserved polyA(+) and polyA(−) motifs provides insights into many polyA and polyT sequences in CRs of insect mt genomes, leading to reveal the mt transcription and its regulation in invertebrates. In addition, we provided a high-quality, well-curated and precisely annotated *D. simulans* mt genome (GenBank: MN611461), which should be included into the NCBI RefSeq database to replace the current reference genome NC_005781.

## Introduction

The annotation of animal mitochondrial (mt) genomes is pivotal to studies on the molecular phylogenetics and evolution of animals, as well as in investigations focusing on RNA processing, maturation, degradation and the regulation of gene expression [1]. In our previous studies, we adopted data from a diverse range of sequencing techniques, including RNA-seq, sRNA-seq, PARE-seq, CAGE-seq, PRO-seq, PA-seq, GRO-seq, PacBio cDNA-seq, Nanopore cDNA-seq, and Nanopore RNA-seq to improve gene annotation, which was defined as pan RNA-seq analysis [2]. By pan RNA-seq analysis, mt genome annotation down to a resolution of 1-bp is achievable, a degree of precision that can be referred to as “precise annotation” [3]. On the basis of these advances, we then performed precise annotation of human, chimpanzee, rhesus macaque, mouse [1], *Erthesina fullo* Thunberg [4] and tick [5] mt genomes, which has led to a number of novel discoveries, notably those with respect to mt non-coding genes (MDL1, MDL1AS, MDL2 and MDL2AS) and the uninterrupted transcription of human mt genome [6].

Precise annotation of transcription initiation sites (TISs) and transcription termination sites (TTSs) sites in the control regions (CRs) of human, chimpanzee, rhesus, and macaque mt genomes has determined that there is only a single TIS_H_ and single TIS_L_ for the heavy strand (H-strand) and light strand (L-strand), respectively, in a mammal mt genome that contains a single CR. On the basis of these annotations, we proposed a novel mt transcription model to describe mammalian mtDNA transcription and its regulation [1]. However, although the so-called “rules” of mtDNA transcription and its regulation may be applicable to vertebrate mt genomes, these established characteristics may not be equally apposite when we attempt to determine TISs and TTSs in the mt genomes of invertebrates, particularly insects, for which patterns tend to be somewhat more complex. Specifically, the mt genomes of many invertebrates contain more than one CR, and these regions are often characterized by a considerably large number of low complexity and short tandem repeat (STR) regions. Moreover, invertebrate CRs typically have considerably low GC contents.

As a CR contains at least one replication origin and two TISs of the mt genome, it is of great importance. The GC contents of CRs in insect mt genomes are much lower than those in mammal mt genomes. For example, the CR in *Drosophila melanogaster* mt genome (RefSeq: NC_024511.2) has a GC content of just 4.28%, notably lower compared to the 46.61% GC content of human mt genome (RefSeq: NC_012920.1). Therefore, the CRs in insect mt genomes are often annotated as AT-rich regions, which contain many polyadenine (polyA) and polythymine (polyT) and AT-rich STRs (e.g., ATATAT) or simple complexity sequences (e.g., ATTAATAT). Until now, only a few motifs or elements in mt CRs have been characterized. In our previous study [1], the H-strand promoters (HSPs), the L-strand promoters (LSPs) and five Conserved Sequence Blocks (CSB1, CSB2, CSB3, LSP and HSP) were annotated in the CRs of 52 mammal mt genomes. In another previous study [7], many sequence motifs in respect to genome replication were identified in AT-rich regions from bacteria (mtDNAs originated from a bacterial ancestor [8]). In a recent study [3], we reported that the tandem repeats in the CR of *E. fullo* mitochondrial genome are associated with TISs. However, the formation and functions of these genomic features are still unclear.

In the present study, with the aim of gaining a better understanding of invertebrate mt transcription and its regulation, we performed precise annotation of the mt genomes of *D. melanogaster*, *D. simulans*, *D. grimshawi*, and *Bactrocera oleae*. Specifically, we sought to: 1) precisely annotate TISs and TTSs in *Drosophila* mt genomes; and (2) determine whether uninterrupted transcription of the H- or L-strand occurs in these genomes, as it does in humans; and (3) provide insights into the formation of the mt AT-rich regions in the light of their functions.

## Results

### Precise annotation of *Drosophila* mitochondrial genomes

Given that PacBio cDNA-seq, Nanopore cDNA-seq, and Nanopore RNA-seq can be used to obtain full-length transcripts [2], we used all available corresponding datasets to facilitate precise annotation of the complete mt genomes of *D. melanogaster* (RefSeq: NC_024511.2), *D. simulans* (RefSeq: NC_005781.1 and GenBank: CM016414.1), and *D. grimshawi* (GenBank: BK006341.1) (**Materials and Methods**). In addition, for comparative purposes, we also precisely annotated the complete mt genome of *B. oleae* (RefSeq: NC_005333.1), a species for which a large high-depth Nanopore cDNA-seq dataset is available online. These new annotations cover both entire strands of mt genomes without gaps or overlaps at a resolution of 1 bp. For each mt genome, the new annotations of two rRNAs (12S rRNA and 16S rRNA), 11 mRNAs encoding 13 proteins, 22 tRNAs and all the antisense genes are presented in **Table 1** for *D. melanogaster* and **Table S2 to S5** (**Supplementary file 1**) for *D. simulans* (RefSeq: NC_005781.1 and CM016414.1), *D. grimshawi*, and *B. oleae*, respectively.

**Table1.**
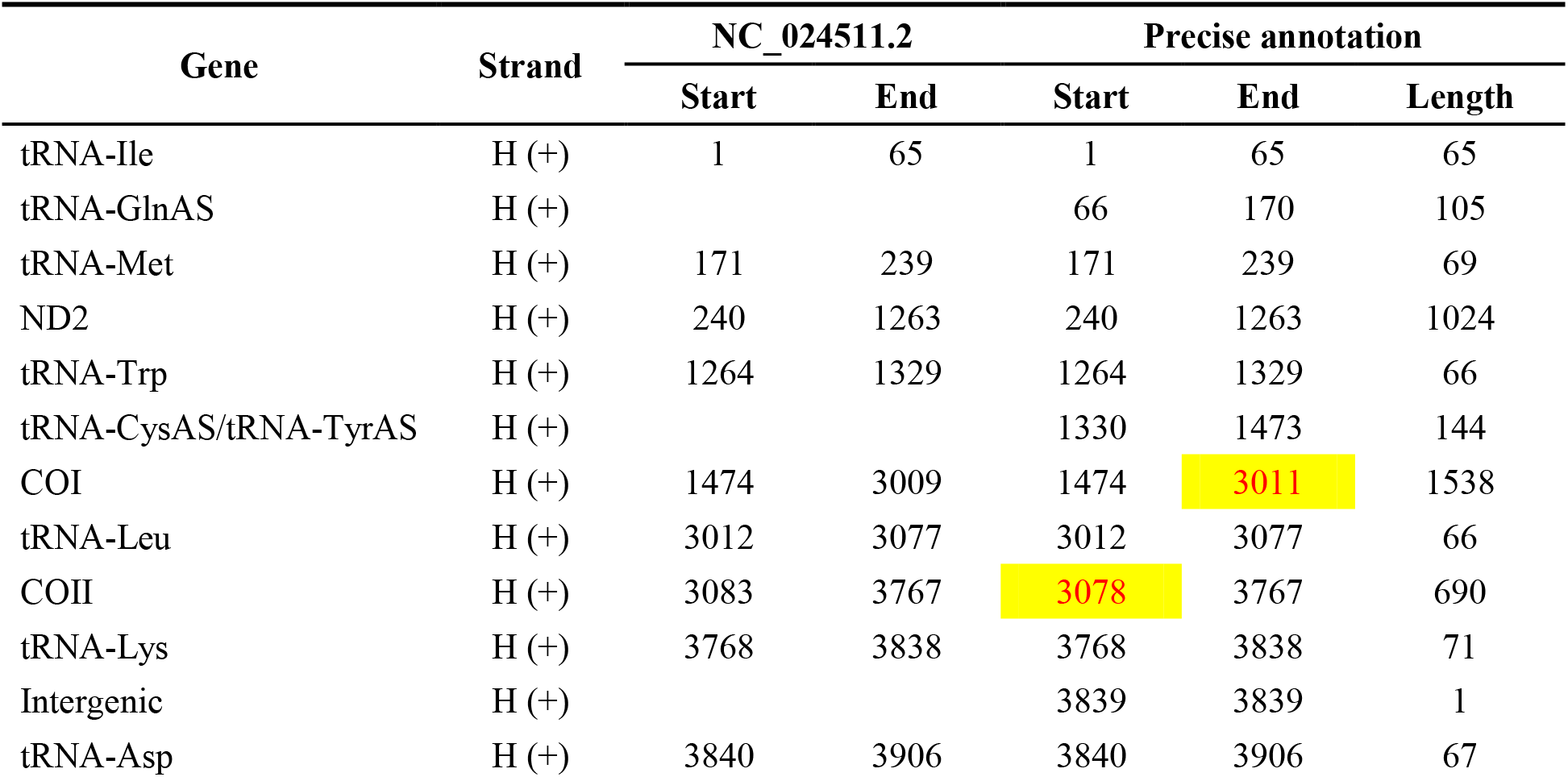
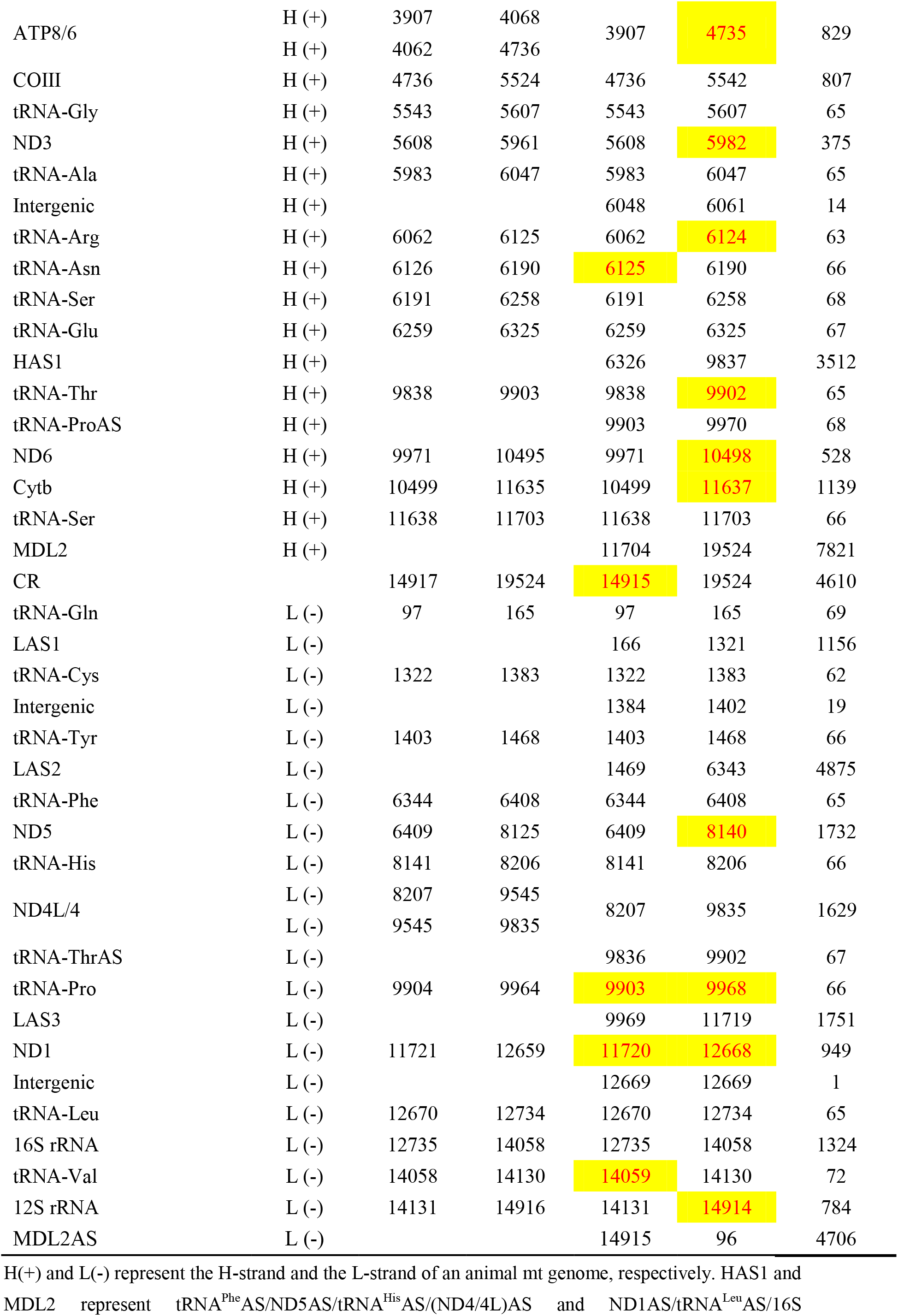
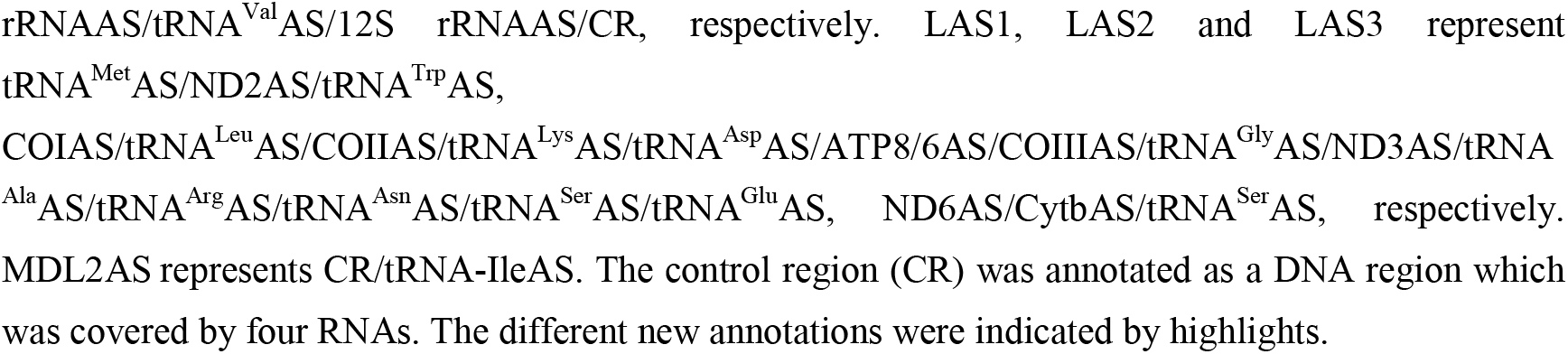
Precise annotation of *Drosophila melanogaster* mt genome.

| Gene | Strand | NC_024511.2 |  | Precise annotation |  |  |
| --- | --- | --- | --- | --- | --- | --- |
|  |  | Start | End | Start | End | Length |
| tRNA-Ile | H (+) | 1 | 65 | 1 | 65 | 65 |
| tRNA-GlnAS | H (+) |  |  | 66 | 170 | 105 |
| tRNA-Met | H (+) | 171 | 239 | 171 | 239 | 69 |
| ND2 | H (+) | 240 | 1263 | 240 | 1263 | 1024 |
| tRNA-Trp | H (+) | 1264 | 1329 | 1264 | 1329 | 66 |
| tRNA-CysAS/tRNA-TyrAS | H (+) |  |  | 1330 | 1473 | 144 |
| COI | H (+) | 1474 | 3009 | 1474 | 3011 | 1538 |
| tRNA-Leu | H (+) | 3012 | 3077 | 3012 | 3077 | 66 |
| COII | H (+) | 3083 | 3767 | 3078 | 3767 | 690 |
| tRNA-Lys | H (+) | 3768 | 3838 | 3768 | 3838 | 71 |
| Intergenic | H (+) |  |  | 3839 | 3839 | 1 |
| tRNA-Asp | H (+) | 3840 | 3906 | 3840 | 3906 | 67 |
| ATP8/6 | H (+) | 3907 | 4068 |  |  |  |
|  | H (+) | 4062 | 4736 | 3907 | 4735 | 829 |
| COIII | H (+) | 4736 | 5524 | 4736 | 5542 | 807 |
| tRNA-Gly | H (+) | 5543 | 5607 | 5543 | 5607 | 65 |
| ND3 | H (+) | 5608 | 5961 | 5608 | 5982 | 375 |
| tRNA-Ala | H (+) | 5983 | 6047 | 5983 | 6047 | 65 |
| Intergenic | H (+) |  |  | 6048 | 6061 | 14 |
| tRNA-Arg | H (+) | 6062 | 6125 | 6062 | 6124 | 63 |
| tRNA-Asn | H (+) | 6126 | 6190 | 6125 | 6190 | 66 |
| tRNA-Ser | H (+) | 6191 | 6258 | 6191 | 6258 | 68 |
| tRNA-Glu | H (+) | 6259 | 6325 | 6259 | 6325 | 67 |
| HAS1 | H (+) |  |  | 6326 | 9837 | 3512 |
| tRNA-Thr | H (+) | 9838 | 9903 | 9838 | 9902 | 65 |
| tRNA-ProAS | H (+) |  |  | 9903 | 9970 | 68 |
| ND6 | H (+) | 9971 | 10495 | 9971 | 10498 | 528 |
| Cytb | H (+) | 10499 | 11635 | 10499 | 11637 | 1139 |
| tRNA-Ser | H (+) | 11638 | 11703 | 11638 | 11703 | 66 |
| MDL2 | H (+) |  |  | 11704 | 19524 | 7821 |
| CR |  | 14917 | 19524 | 14915 | 19524 | 4610 |
| tRNA-Gln | L (-) | 97 | 165 | 97 | 165 | 69 |
| LAS1 | L (-) |  |  | 166 | 1321 | 1156 |
| tRNA-Cys | L (-) | 1322 | 1383 | 1322 | 1383 | 62 |
| Intergenic | L (-) |  |  | 1384 | 1402 | 19 |
| tRNA-Tyr | L (-) | 1403 | 1468 | 1403 | 1468 | 66 |
| LAS2 | L (-) |  |  | 1469 | 6343 | 4875 |
| tRNA-Phe | L (-) | 6344 | 6408 | 6344 | 6408 | 65 |
| ND5 | L (-) | 6409 | 8125 | 6409 | 8140 | 1732 |
| tRNA-His | L (-) | 8141 | 8206 | 8141 | 8206 | 66 |
| ND4L/4 | L (-) | 8207 | 9545 | 8207 | 9835 | 1629 |
|  | L (-) | 9545 | 9835 |  |  |  |
| tRNA-ThrAS | L (-) |  |  | 9836 | 9902 | 67 |
| tRNA-Pro | L (-) | 9904 | 9964 | 9903 | 9968 | 66 |
| LAS3 | L (-) |  |  | 9969 | 11719 | 1751 |
| ND1 | L (-) | 11721 | 12659 | 11720 | 12668 | 949 |
| Intergenic | L (-) |  |  | 12669 | 12669 | 1 |
| tRNA-Leu | L (-) | 12670 | 12734 | 12670 | 12734 | 65 |
| 16S rRNA | L (-) | 12735 | 14058 | 12735 | 14058 | 1324 |
| tRNA-Val | L (-) | 14058 | 14130 | 14059 | 14130 | 72 |
| 12S rRNA | L (-) | 14131 | 14916 | 14131 | 14914 | 784 |
| MDL2AS | L (-) |  |  | 14915 | 96 | 4706 |

H(+) and L(-) represent the H-strand and the L-strand of an animal mt genome, respectively. HAS1 and
MDL2 represent tRNA<sup>Phe</sup>AS/ND5AS/tRNA<sup>His</sup>AS/(ND4/4L)AS and ND1AS/tRNA<sup>Leu</sup>AS/16S
rRNAAS/tRNA<sup>Val</sup>AS/12S rRNAAS/CR, respectively. LAS1, LAS2 and LAS3 represent tRNA<sup>Met</sup>AS/ND2AS/tRNA<sup>Trp</sup>AS, COIAS/tRNA<sup>Leu</sup>AS/COIIAS/tRNA<sup>Lys</sup>AS/tRNA<sup>Asp</sup>AS/ATP8/6AS/COIIIAS/tRNA<sup>Gly</sup>AS/ND3AS/tRNA<sup>Ala</sup>AS/tRNA<sup>Arg</sup>AS/tRNA<sup>Asn</sup>AS/tRNA<sup>Ser</sup>AS/tRNA<sup>Glu</sup>AS, ND6AS/CytbAS/tRNA<sup>Ser</sup>AS, respectively. MDL2AS represents CR/tRNA-IleAS. The control region (CR) was annotated as a DNA region which was covered by four RNAs. The different new annotations were indicated by highlights.

Although a majority of the new annotations (**Table 1 and Table S2-S5**) are consistent with the previous annotations of the mt genomes archived in the NCBI RefSeq or GenBank databases, a few new annotations were used to correct or modify the previous annotations, representing significant improvements. Here, as an illustrative example of these improved annotations, we describe the precise annotation of *D. melanogaster* mt genome (**Table 1**). The annotations of five tRNA genes (tRNA^Arg^, tRNA^Asn^, tRNA^Thr^, tRNA^Pro^ and tRNA^Val^) were corrected by precise annotation, since the sequences of these tRNAs according to the previous annotations were insufficiently complete to enable predictions of tRNA secondary structures (**Figure 1A**). The annotations of eight mRNA genes (COI, COII, ATP8/6, ND3, ND6, Cytb, ND5, and ND1) were corrected or modified such that the mRNA sequences are longer than the corresponding coding sequences (CDSs). For example, the annotation of COI was modified from 1474-30<u>**09**</u> to 1474-30<u>**11**</u>. The annotation of ND1 was corrected from 1172<u>**1**</u>-126<u>**59**</u> to 1172<u>**0**</u>-126<u>**68**</u> by adding nine nucleotide residues (<u>**TTG**</u>TTTTAT) to the 5ʹ end of the previously annotated ND1 sequence. As a consequence of this correction, the start codon of ND1 is now predicted to be TTG, rather than ATA, which was assumed according to the previous annotation (RefSeq: NC_024511.2). This correction confirmed that the CDS regions of mt coding genes can not be annotated accurately based on analysis of the predicted open reading frames (ORFs) [3]. A further important amendment is the modification of the CR annotation from 1491<u>**7**</u>–19524 to 1491<u>**5**</u>–19524 (**Table 1**), and subsequently, the annotation of 12S rRNA was modified from 14131–1491<u>**6**</u> to 14131–1491<u>**4**</u>.

**Figure 1.**
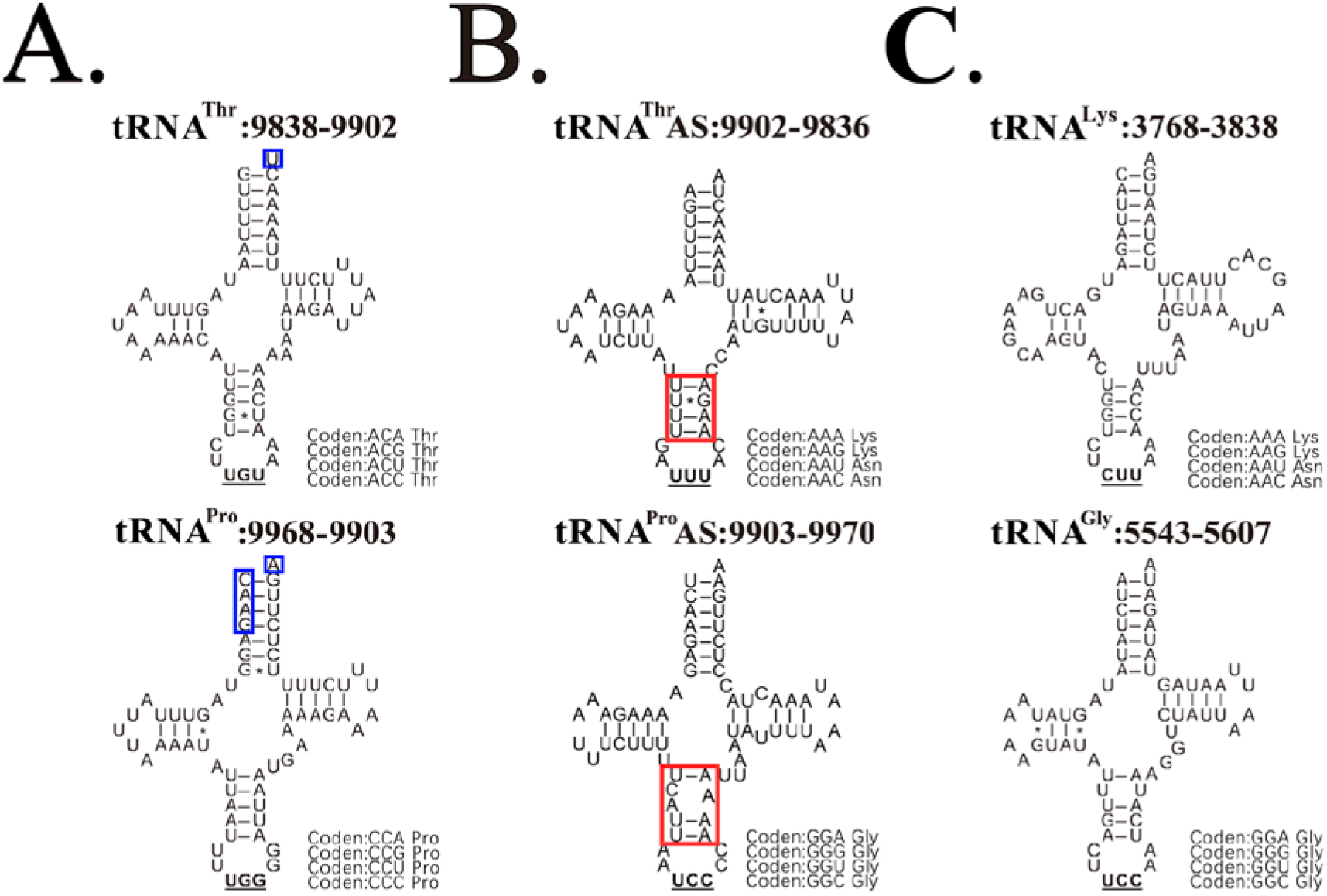
Two antisense tRNAs with CCAs at the 3’ ends. All the genomic positions refer to the *Drosophila melanogaster* mt genome (RefSeq: NC_024511.2). **A**. The annotations of tRNA^Thr^ and tRNA^Pro^ were corrected by precise annotation. The sequences of these tRNAs according to the previous annotations (incomplete nucleotide residues marked in blue boxes) were not complete to form tRNA secondary structures. **B**. Two antisense tRNAs (tRNA^Thr^AS and tRNA^Pro^AS) have CCAs at their 3’ ends. The anticodon arm (marked in red boxes) of tRNA^Pro^AS is unlikely to be stable for normal function. **C**. tRNA^Thr^AS and tRNA^Pro^AS may function as tRNA^Lys^ and tRNA^Gly^.

### A reference sequence of *Drosophila simulans* mt genome

The mt genomes of *D. simulans* (RefSeq: NC_005781.1 and GenBank: CM016414.1) were precisely annotated using PacBio cDNA-seq (ENA: ERP110987). However, the mt genome NC_005781.1 does not contain the sequence of CR and the mt genome CM016414.1 has several assembly errors. Then, we had to assembly a new reference sequence of *D. simulans* mt genome and performed precise annotation of it. The new reference mt genome for *D. simulans* (**Supplementary file 2**) has already been submitted to the NCBI GenBank database under the accession number MN611461 with precise annotations (**Table S6**). On the basis of a comparison among the three *D. simulans* mt genomes (RefSeq: NC_005781.1, GenBank: CM016414.1, and MN611461), we were able to determine the aforementioned errors in the CM016414.1 sequence, the most significant of which is that the final 201 bp of the COII sequence has been incorrectly duplicated (**Table S5**). A further prominent error is that of a 1-bp deletion in a ployA region (CM016414: 8628-8637) of ND1, which would have resulted in an incorrectly translated protein. Accordingly, given our relevant findings—(1) the current reference genome NC_005781.1 does not contain a CR sequence; (2) the mt genome CM016414.1 has several assembly errors; and (3) the high-quality, well-curated, and precisely annotated MN611461 has very high nucleotide identities of 97.15% and 99.9% with NC_005781.1 and CM016414.1, respectively—we believe that the newly annotated *D. simulans* mt genome (GenBank: MN611461) should replace the NC_005781.1 reference sequence in the NCBI RefSeq database.

A comparison of the new annotations of all *Drosophila* and *B. oleae* mt genomes enabled us to determine the start codons of all mRNA genes. Notable in this respect are the CCG, ATC, and GTG start codons of COI, ATP8/6 and ND5, respectively, in *D. simulans* as these are seldom used as start codons in insect mt genomes and were not identified in previous precise annotations of the *Erthesina fullo* [3] and tick [5] mt genomes. In this regard, we incorrectly reported the start codon of COI in the *D. silvarum* mt genome (GenBank: MN347015) as ATA in our previous study [5], which needs to be corrected to CCG. The new annotations reveal that the ORFs of mRNAs in *Drosophila* and *B. oleae* mt genomes starts at their first positions with the exception of COII, while the ORFs of COII start at their 5^th^ or 6^th^ positions, as there are several nucleotide residues (AATAA in *D. melanogaster*) prior to the start codon of ATG. Although the intergenic regions in *Drosophila* mt genomes were cleaved between neighboring tRNAs to form small RNAs (sRNAs), these are unlikely to have biological functions, in our view. The longest intergenic sequence we detected is a 28-nt region (ATATACATATACCTCTAATAATTAAATA) between tRNA^Arg^ and tRNA^Asn^ in *B. oleae* mt genome, thereby confirming that apart from a novel 31-nt ncRNA, the intergenic regions between mt tRNA genes in insects are longer than those in mammals [5]. One of two important findings in the present study is that two antisense tRNAs (tRNA^Thr^AS and tRNA^Pro^AS) in the *D. melanogaster* mt genome (RefSeq: NC_024511.2) have CCAs at their 3ʹ ends (**Figure 1B**), which differs from other antisense tRNAs that are characterized by polyA 3ʹ terminal sequences. Although these two antisense tRNAs may function as tRNA^Lys^ and tRNA^Gly^ (**Figure 1C**), respectively, the anticodon arm of tRNA^Pro^AS is unlikely to be stable for normal function. Further investigations are thus required to confirm a previously proposed hypothesis, which states that antisense tRNAs may function as tRNAs [9].

### Discovery of polyA motifs in insect mt genomes

In a series of previous studies, we preformed precise annotation of human, chimpanzee, rhesus macaque, mouse [1], *Erthesina fullo* Thunberg [3], and tick [5] mt genomes and thereby provided novel insights into the mammalian mtDNA transcription and its regulation. On the basis of these findings, we proposed an “mt transcription” model [3], in which we envisage that a single CR initiates transcription of the H- and L-strands of mammal mt genomes at the TIS_H_ and TIS_L_ sites, respectively, and that in response to the binding of mitochondrial transcription termination factors, multiple TTSs function as switches to determine the production of short or long primary transcripts or uninterrupted transcription, rather than completely terminating mt transcription. In a mammal mt genome, a CR is involved in the synthesis of four RNAs (**Figure 2A**), namely, two on the H-strand, the shorter mt D-loop 1 (MDL1) and longer MDL2, and two on L-strand, the shorter mt D-loop 1 antisense gene (MDL1AS) and the longer MDL2AS [6]. Using PacBio cDNA-seq data from *D. simulans*, we precisely annotated the TIS_H_ and the TIS_L_ in *Drosophila simulans* mt genome and thereby found that they are located in polyA motifs close to the 5ʹ and 3ʹ ends of the CR, which are defined as polyA(+) and polyA(−) motifs, respectively, and include all adenine residues and 10 downstream nucleotide residues (**Figure 2B**). However, we were unable to detect either TIS_H_ or TIS_L_ in the polyA(+) and polyA(−) motifs of *D. melanogaster*, *D. grimshawi*, or *B. oleae* mt genomes, using Nanopore cDNA-seq data. Conversely, we identified several TIS_H_ and TIS_L_ in the CR of *B. oleae* mt genome (**Figure 2C**), although owing to the general low quality of the Nanopore cDNA-seq data, these sites will need further confirmation. Given that these polyA(+) and polyA(−) motifs in animal mt genomes need to be recognized by the same transcription system, they tend to be well conserved, and accordingly have a high degree of nucleotide identity. Further analysis revealed that the polyA(+) and polyA(−) motifs in CRs (**Table 2**) are conserved in the *Drosophila* genus and numerous other insect genera (e.g., *Procecidochares utilis*). We had previously established that polyA motifs in the CRs of mammal mt genomes are also associated with mt TISs, although the lengths of the polyA sequences are typically very short (e.g., 3 bp in the human mt genome). These findings suggested that the polyA(+) and polyA(−) motifs in CRs may function as signals to initiate mtDNA transcription.

**Figure 2.**
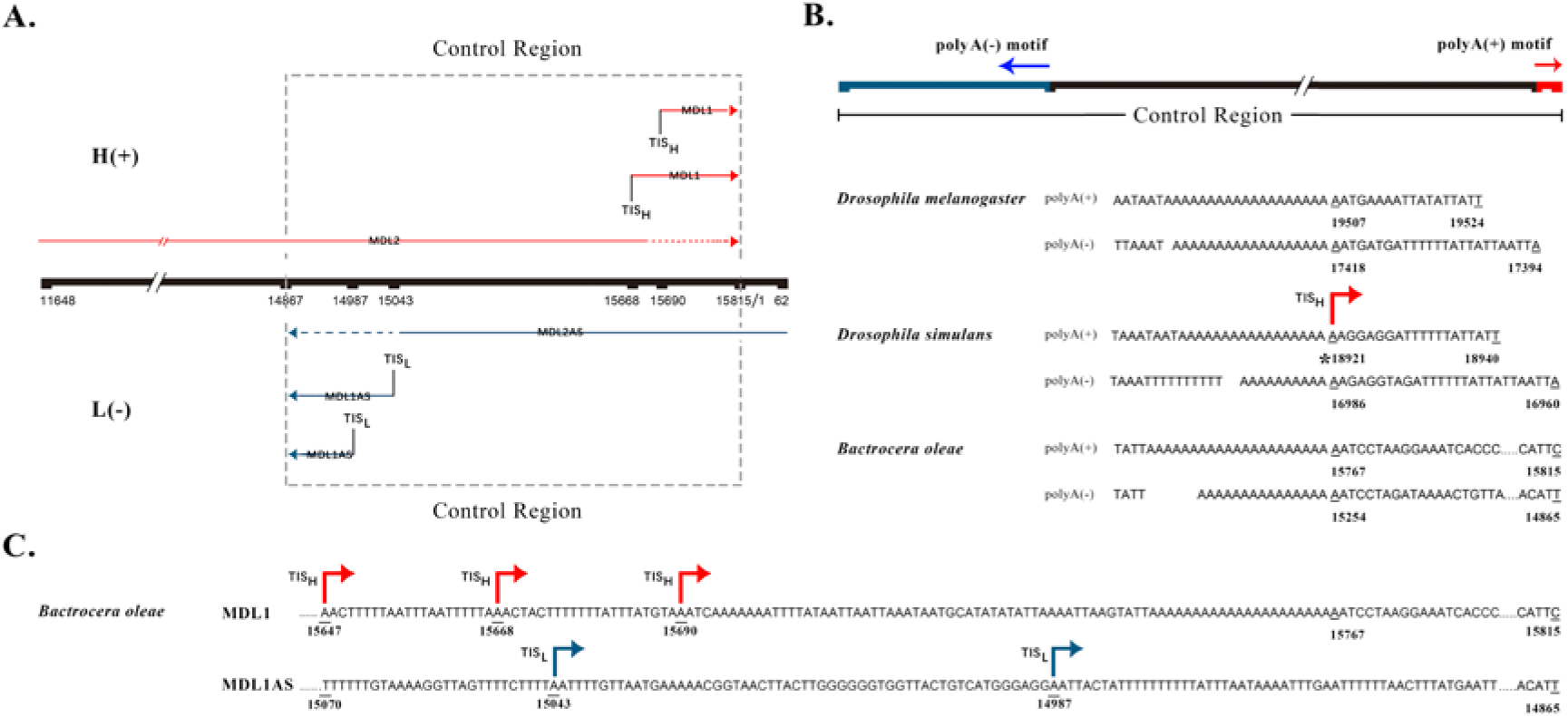
Annotation of polyA(+) and polyA(−) motifs in control regions. **A**. One control region initiates the transcription of the H-strand and the L-strand of a mammal mt genome at the TIS_H_ and the TIS_L_, respectively. However, the “rules” are unlikely to be longer applicable in invertebrate mt genomes, as the situation is rather complicated. Although the positions of MDL1, MDL1AS, MDL2, MDL2AS, TIS_H_ and the TIS_L_ were annotated in *Bactrocera oleae* mt genome (RefSeq: NC_005333.1), these annotations still needs the final determination. **B**. The locations of polyA(+) and polyA(−) motifs in control regions of insect mt genomes, particularly in *Drosophila melanogaster* (RefSeq: NC_024511.2), *D. simulans* (GenBank: CM016414.1) and *Bactrocera oleae* (RefSeq: NC_005333.1) mt genomes. Among these polyA(+) and polyA(−) motifs, only the polyA(+) motif of *D. simulans* were detected to be associated with TIS_H_. **C**. Several TIS_H_ and TIS_L_ were detected in the CR of *B. oleae* mt genome using Nanopore cDNA-seq data. However, they still needs the final determination.

**Table2.** PolyA motifs in insect mt genomes.

| Species | Accession<br>Number | CR range | GC(%) | Start |
| --- | --- | --- | --- | --- |
| *Drosophila melanogaster* | NC_024511.2 | 14917-19524 | 4.28 | PolyA(+):19488<br>PolyA(-):17436 |
| *Drosophila simulans* | CM016414.1 | 11401-15497 | 6.71 | PolyA(+):15365<br>PolyA(-):13457 |
| *Bactrocera oleae* | NC_005333.1 | 14867-15815 | 13.08 | PolyA(+):15746<br>PolyA(-):15269 |
| *Drosophila mercatorum* | NC_044669.1 | 15006-16024 | 8.25 | PolyA(+):15987<br>PolyA(-):15538 |
| *Drosophila formosana* | NC_028518.1 | 15015-16100 | 7.56 | PolyA(+):16059<br>PolyA(-):15590 |
| *Drosophila incompta* | NC_025936.1 | 15514-606 | 8.61 | PolyA(+):563<br>PolyA(-):118 |
| *Drosophila littoralis* | NC_011596.1 | 14995-16017 | 9.88 | PolyA(+):15976<br>PolyA(-):15524 |
| *Drosophila yakuba* | NC_001322.1 | 14943-16019 | 7.16 | PolyA(+):15983<br>PolyA(-):15592 |
| *Drosophila santomea* | NC_023825.1 | 14945-16022 | 7.15 | PolyA(+):15984<br>PolyA(-):15596 |
| *Procecidochares utilis* | NC_020463.1 | 14995-15922 | 9.48 | PolyA(+):15854<br>PolyA(-):15685 |

The “start” column indicates the genomic position of the first nucleotide residue of a polyA(+) or a polyA(-) motif. The range of control region (CR) was annotated by new annotations, if the previous annotations were corrected or modified.

## Conclusion and Discussion

In the present study, we performed precise annotation of *D. melanogaster*, *D. simulans*, *D. grimshawi*, *Bactrocera oleae* by pan RNA-seq analysis. Two important findings were reported for the first time, including the discovery of the conserved polyA(+) and polyA(−) motifs in the CRs and the adding of CCAs to the 3’ ends of two antisense tRNAs. Based on the above findings, we proposed: (1) polyA/polyT in CRs may function as signals to initiate mtDNA transcription; (2) the duplication, recombination or mutation of these polyA/polyT sequences formed the AT-region regions during evolution; and (3) since CRs of many invertebrate species still contain many polyA/polyT sequences, there is a high probability that several TISs and TTSs exist in invertebrate mt genomes. The key step leading to the proposal of our model was that the conserved polyA(+) and polyA(−) motifs in the mt genomes of the *Drosophila* genus have more than 10 adenine residues, significantly longer than those in many other insect genera on average. The discovery of the conserved polyA(+) and polyA(−) motifs provides insights into many polyA/polyT sequences in CRs of insect mt genomes, leading to reveal the mt transcription and its regulation in invertebrates.

## Materials and Methods

Five complete mt genomes of *Drosophila melanogaster* (RefSeq: NC_024511.2), *D. simulans* (RefSeq: NC_005781.1 and GenBank: CM016414.1), *D. grimshawi* (GenBank: BK006341.1) and *Bactrocera oleae* (RefSeq: NC_005333.1) were downloaded from the NCBI RefSeq or GenBank databases. Five small RNA-seq (sRNA-seq) datasets (SRP017617, SRP199841, ERP011832, ERP012885 and ERP106913) were used to annotate the *D. melanogaster* mt genome The datasets SRP302633 (Pacbio cDNA-seq) of *D. melanogaster*, ERP117816 (Nanopore cDNA-seq) of *D. melanogaster*, ERP110987 (Pacbio cDNA-seq) of *D. simulans*, SRP135764 (Pacbio cDNA-seq) of *D. grimshawi* and SRP189010 (Nanopore cDNA-seq) of *B. oleae* were used for precise annotation. All the sRNA-seq, Pacbio cDNA-seq and Nanopore cDNA-seq datasets were downloaded from the NCBI SRA or EBI ENA databases.

Precise annotation of mt genomes followed the protocol published in one of our previous study [1]. Data cleaning and quality control were performed using Fastq_clean [10]. The PacBio IsoSeq3 was utilized to transform subreads to CCSs (circular consensus sequences) with parameters (Minimum Full Passes 1), and then remove primers, concatemers and ployA tails with default parameters for further analysis. Statistics and plotting were conducted using the software R v2.15.3 with the package ggplot2 [11]. The 5’ and 3’ ends of mature transcripts, polycistronic transcripts, and antisense transcripts were observed and curated using the software Tablet v1.15.09.01 [12]. Each identified transcript was required to be supported by at least three CCSs (**Supplementary file 3**).

## Supplementary information

### Declarations

#### Ethics approval and consent to participate

Not applicable.

#### Consent to publish

Not applicable.

#### Availability of data and materials

The *Drosophila simulans* mitochondrial genome is available at the NCBI GenBank database under the accession numbers MN611461, which was also provided in supplementary file 2.

#### Authors’ contributions

Shan Gao conceived the project. Shan Gao and Wenjun Bu supervised this study. Guangcai Liang performed programming. Jia Chang, Bingjun He, Tung On Yau and Xin Li downloaded, managed and processed the data. Shan Gao drafted the main manuscript text. Shan Gao and Jishou Ruan revised the manuscript.

## Acknowledgments

We are grateful for the help from the following faculty members of College of Life Sciences at Nankai University: Tao Zhang, Dawei Huang, Qiang Zhao, Huaijun Xue and Zhen Ye. We would like to thank Editage (www.editage.cn) for polishing part of this manuscript in English language. This manuscript was online as a preprint on Feb 5^th^, 2021 at xx.

